# Phasic norepinephrine is a neural interrupt signal for unexpected events in rapidly unfolding sensory sequences – evidence from pupillometry

**DOI:** 10.1101/466367

**Authors:** Sijia Zhao, Maria Chait, Fred Dick, Peter Dayan, Shigeto Furukawa, Hsin-I Liao

**Author notes:** Corresponding Authors: Maria Chait, Ear Institute, University College London, 332 Gray’s Inn Road, London WC1X 8EE, UK; Sijia Zhao, Ear Institute, University College London, 332 Gray’s Inn Road, London WC1X 8EE, UK. Lead Contact: Maria Chait, Ear Institute, University College London, 332 Gray’s Inn Road, London WC1X 8EE, UK.

## Abstract

The ability to track the statistics of our surroundings is a key computational challenge. A prominent theory (Dayan & Yu, 2006) proposes that the brain monitors for ‘unexpected uncertainty’ – events which deviate substantially from model predictions, indicating model failure. Norepinephrine (NE) is thought to play a key role in this process by serving as an interrupt signal, initiating model-resetting. However, evidence is from paradigms where participants actively monitored stimulus statistics. To determine whether NE routinely reports the statistical structure of our surroundings, even when not behaviourally relevant, we used rapid tone-pip sequences that contained perceptually salient pattern-changes associated with abrupt structural violations vs. emergence of regular structure. Phasic pupil dilations (PDR) were monitored to assess NE. We reveal a remarkable specificity: When not behaviourally relevant, only abrupt structural violations evoked a PDR. The results demonstrate that NE tracks ‘unexpected uncertainty’ on rapid time scales relevant to sensory signals.

## Introduction

A growing body of work demonstrates that observers maintain detailed models of the statistics of their environments over various timescales, combining this information with sensory input to inform choice (Rushworth and Behrens, 2008), increase response accuracy (Krishnamurthy et al., 2017), speed up reaction times (Bestmann et al., 2008; Marshall et al., 2016), and improve detection (Sohoglu and Chait, 2016; Southwell and Chait, 2018). A key challenge in this context is keeping track of the evolving input statistics so as to ensure model validity. Here we investigated automatic and controlled aspects of the neural response to this challenge in a fast-paced domain.

For effective model maintenance, a central dilemma faced by the brain is to arbitrate between gradual and punctate changes in the environment (Gershman et al., 2013). In the former case, model-updating progresses at a steady pace, dictated by the model’s estimate of local noise (‘expected uncertainty’) arising from tracked environmental stochasticities (Bland and Schaefer, 2012; O’Reilly, 2013). However, environments can also change substantially and suddenly. The ability to detect such change points is crucial for optimal behavior, because they indicate that the observer’s beliefs about the environment are no longer a valid representation of reality, and should be reset (Marshall et al., 2016; Payzan-LeNestour et al., 2013). For example, in Nassar et al. (2012) subjects were instructed to predict sequentially-presented numbers drawn from a Gaussian distribution whose mean occasionally changed abruptly. Following change points, participants tended to alter their behavior in a way that reflected abandonment of old expectations and more rapid acquisition of new ones - e.g., recent events had more influence on decisions than those occurring further in the past, equivalent to an increased learning (and forgetting) rate.

Although such change processes can often be described optimally in hierarchical probabilistic terms, an alternative heuristic is for the brain to monitor events that fall outside the threshold of ‘expected uncertainty’ estimated for the model, and treat them as signaling potential change points in the environment. Such so-called ‘unexpected uncertainty’ (Dayan and Yu, 2006) has been suggested as interrupting top-down processes so as to prioritize bottom-up evidence accumulation, thereby speeding up discovery of the new structure of the environment (Dayan and Yu, 2006; O’Reilly, 2013; Sara and Bouret, 2012). The neuromodulator norepinephrine (alternatively noradrenaline or NE) has been hypothesized to play a critical role in this updating process (Bouret and Sara, 2005; Dayan and Yu, 2006; Marshall et al., 2016; Payzan-LeNestour et al., 2013; Yu and Dayan, 2005). NE is generated in the brainstem nucleus Locus Coeruleus (LC), which projects extensively across the brain and spinal cord (Samuels and Szabadi, 2008a, 2008b; Sara and Bouret, 2012) and is thus optimally placed to signal a global state change in the environment. However, a vast literature has also implicated NE in controlling vigilance, orienting behavior, selective attention and surprise (Aston-Jones and Cohen, 2005; Avery and Krichmar, 2017; Sara and Bouret, 2012), suggesting that it might instead play a much less specific role, associated with regulating arousal.

The bulk of work on NE and model updating in humans has involved paradigms in which participants actively monitor the statistics of the stimulus, for instance through explicit tracking tasks (Krishnamurthy et al., 2017; Nassar et al., 2012; Payzan-LeNestour et al., 2013; Preuschoff et al., 2011) or in speeded stimulus-response paradigms (Marshall et al., 2016). It is therefore an open question whether NE involvement is driven by behavioral relevance (for decisions or motor responses), or if NE plays a more ubiquitous role in reporting changes in the statistical structure of our surroundings. In the latter case, we need to examine which events trigger its release.

Sensory systems continuously analyze probabilistic information which unfolds on a rapid timescale, even when this information is not immediately relevant to behavior (Barascud et al., 2016; Sohoglu and Chait, 2016; Turk-Browne et al., 2010). It is therefore compelling to ask **(1)** how the fast-paced and automatic mechanisms that detect changes in statistics within rapid sensory signals interface with NE, **(2)** how NE’s involvement compares with other aspects of neural dynamics, and **(3)** what effect there is, if any, of making the changes behaviorally consequential. In addition, by understanding the contingencies to which NE responds, we hope to gain extra clarity on the heuristic separation between gradual and punctate change that is critical for effective model maintenance.

To examine these questions, we sought a sensory paradigm that induces such changes, along with a way of assessing the effect on NE and other neural systems. For the first, we considered rapid auditory patterns (Fig. 1A) consisting of sequences of tone-pips (new on each trial) containing transitions either from a repeating or ‘regular’ (REG) to a random (RAND) frequency structure, or the reverse. These stimuli are particularly suited for our purposes since at a presentation rate of 20Hz, the sequences are too rapid for naive listeners to explicitly follow the unfolding pattern. Rather, the changes in sequence structure (in both directions) readily ‘pop out’ from the stimulus stream irrespective of subjective effort (see stimulus examples in sup. materials or https://goo.gl/vddYuS). Furthermore, the changes induce patterns of neural dynamics (Barascud et al., 2016; Southwell et al., 2017) which hint that they might illuminate the central dilemma about punctate versus gradual change. Transitions from regular to random frequency structures evoke a prompt mismatch neural response, triggered by the abrupt violation of the regular pattern. The opposite transitions, random to regular - despite having matched overall spectro-temporal structure and being similarly detectable - do not generate a mismatch response. Instead, the dynamics of the brain response are consistent with an evidence accumulation process which changes more slowly from one structure to the other.

**Figure 1:**
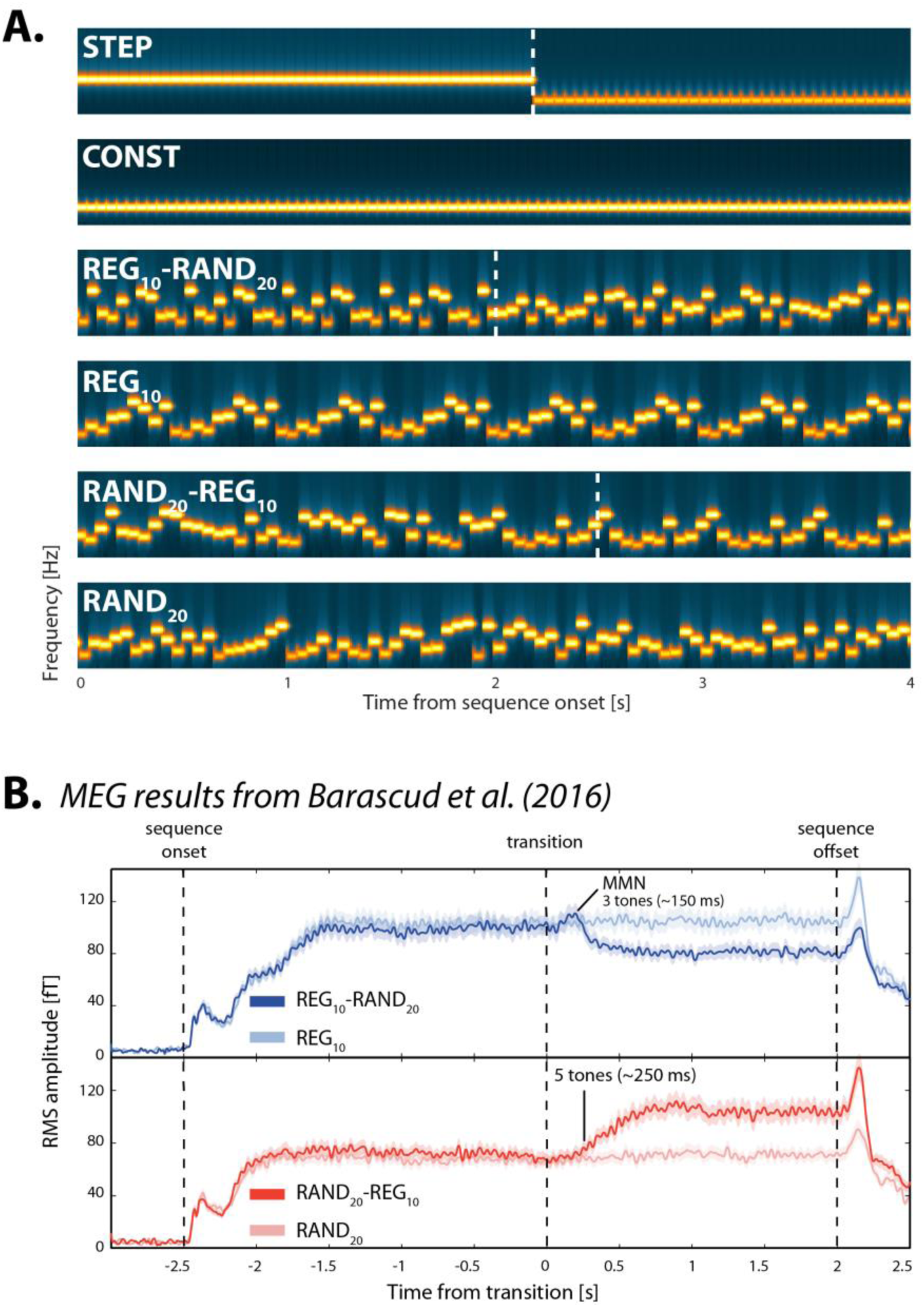
Example spectrograms of basic stimuli, and brain responses to REG-RAND and RAND-REG transitions recorded with MEG. [A] The stimuli were sequences of concatenated tone-pips (50ms) with frequencies drawn from a pool of 20 fixed values. The tone-pips were arranged according to 6 frequency patterns, generated anew for each subject and on each trial: CONST sequences consisted of a single repeating tone; STEP contained a step change from one tone frequency to another; REG_10_ sequences were generated by randomly selecting 10 frequencies from the pool and iterating that sequence to create a regularly repeating pattern; RAND_20_ were generated by randomly sampling from the full pool with replacement; REG_10_-RAND_20_ and RAND_20_-REG_10_ sequences contained a transition between a regular and random pattern or vice versa. Transition times (between 2.5 - 3.5s post onset) are indicated by a white dashed line. In RAND_20_-REG_10_ sequences, the transition time is defined as occurring after the first full regularity cycle, i.e. once the transition becomes statistically detectable. For presentation purposes only, the plotted sequence lengths are equal. Durations varied randomly between 5 to 7s. [B] Brain responses (N=13) to REG_10_-RAND_20_ (top panel) and RAND_20_-REG_10_ (bottom panel), together with their no-change controls, recorded with magnetoencephalography (MEG). Plotted is Root Mean Square (RMS) over channels, as an estimate of instantaneous power. The figures show the entire stimulus epoch, relative to the transition. Shaded areas are ±1 SEM. The transition from RAND to REG is associated with a gradual increase in sustained power from ∼250ms (5 tones) post transition. The transition from REG to RAND evokes an MMN-like response (at ∼150ms after the transition) followed by a sharp drop in the sustained response. These changes in power are hypothesized to reflect the instantiation (RAND-REG) or interruption (REGRAND) of a contextual top-down model. See Barascud et al. (2016) for more details.

In order to assess NE, we turned to the eyes. Indirect measures of NE release can be obtained from monitoring non-luminance-mediated changes in pupil size (Aston-Jones and Cohen, 2005; Joshi et al., 2016). This renders pupillometry an attractive, non-invasive means of probing NE activity in the brain. There is a consistent mechanistic correlation between spiking activity in the LC and changes in pupil size, both when spontaneously occurring, and when triggered by external events (Joshi et al., 2016; Reimer et al., 2016). In particular, transient pupil dilation responses (PDR) have been shown to causally relate to phasic activity within the LC-NE system (Joshi et al., 2016; Reimer et al., 2016), though there remains uncertainty about the specific circuitry (Costa and Rudebeck, 2016). Capitalizing on these links, recent pupillometry studies have revealed a relationship between pupil dilation and predictability (Krishnamurthy et al., 2017; Nassar et al., 2012), providing (indirect) evidence for the involvement of phasic LC-NE responses in signaling uncertainty.

Thus, we monitored pupil size whilst subjects listened to changing auditory sequences, including the disambiguating regular-random (REG-RAND) and random-regular (RAND-REG) transition types. If the pupil-linked LC-NE system generally monitors for salient state changes in the environment, both transition types are expected to evoke PDRs. However, under the hypothesis that phasic LC-NE responses are selective for punctate changes even when detected automatically in speeded inputs, we should observe pupil dilation responses to the former, but not the latter, transition. By also asking participants explicitly to monitor for and report both types of transitions, we could see if the coupling to NE responses is obligatory or behaviorally penetrable – i.e., whether the range of statistical model violations that determine NE activity can be influenced by instructions.

## Results

The basic stimulus set is shown in Fig. 1A. In referring to the stimuli we adopt a nomenclature where the term in uppercase denotes the type of signal (RAND vs REG) and the subscript indicates to the size of the sub-pool from which the relevant pattern is created. Thus, RAND_20_ is a tone series created by randomly selecting each tone (with replacement) from a full pool of 20 frequencies. RAND_10_ is a series created from a subset of 10 different frequencies (randomly selected from the full pool), while REG_10_ is a regular pattern consisting of a repeating sequence of 10 tones (a different pattern, and a different sub-pool on each trial).

Brain response (MEG and EEG) data suggest that while REG_10_-RAND_20_ and RAND_20_-REG_10_ transitions are characterized by opposite statistics (emergence vs. violation of regularity), both are detected automatically, and at a similar latency even when participants’ attention is directed elsewhere (Barascud et al., 2016; Southwell et al., 2017). When asked to respond behaviorally to transitions, listeners exhibit ceiling performance and similar reaction times with comparable variability (Barascud et al. 2016, also replicated here in Exp. 3). Thus, these signals are well suited for disambiguating the role of the pupil linked LC-NE system in tracking statistics of rapidly-evolving sensory signals.

To control for overall engagement (Jepma and Nieuwenhuis, 2011) and ensure broad attention to the auditory stimuli, but <u>without</u> requiring active tracking of the transitions, naïve participants (in Exp. 1,2,4) detected short silent gaps as they listened to the tone-pip sequences. Gap occurrence was uncorrelated to state transition, and the subset of sequences containing gaps were excluded from analyses. (In Exp. 3 we investigate the effect of making the state transition task-relevant).

## Exp. 1: The pupil dilates to violation but not emergence of regularity in complex tone patterns

### Exp. 1A (N=18)

Fig. 2A plots the average normalized pupil size data across all participants as a function of time relative to the transition. Clear PDRs were observed in the STEP and REG_10_-RAND_20_ conditions, but not in the RAND_20_-REG_10_ condition.

**Figure 2:**
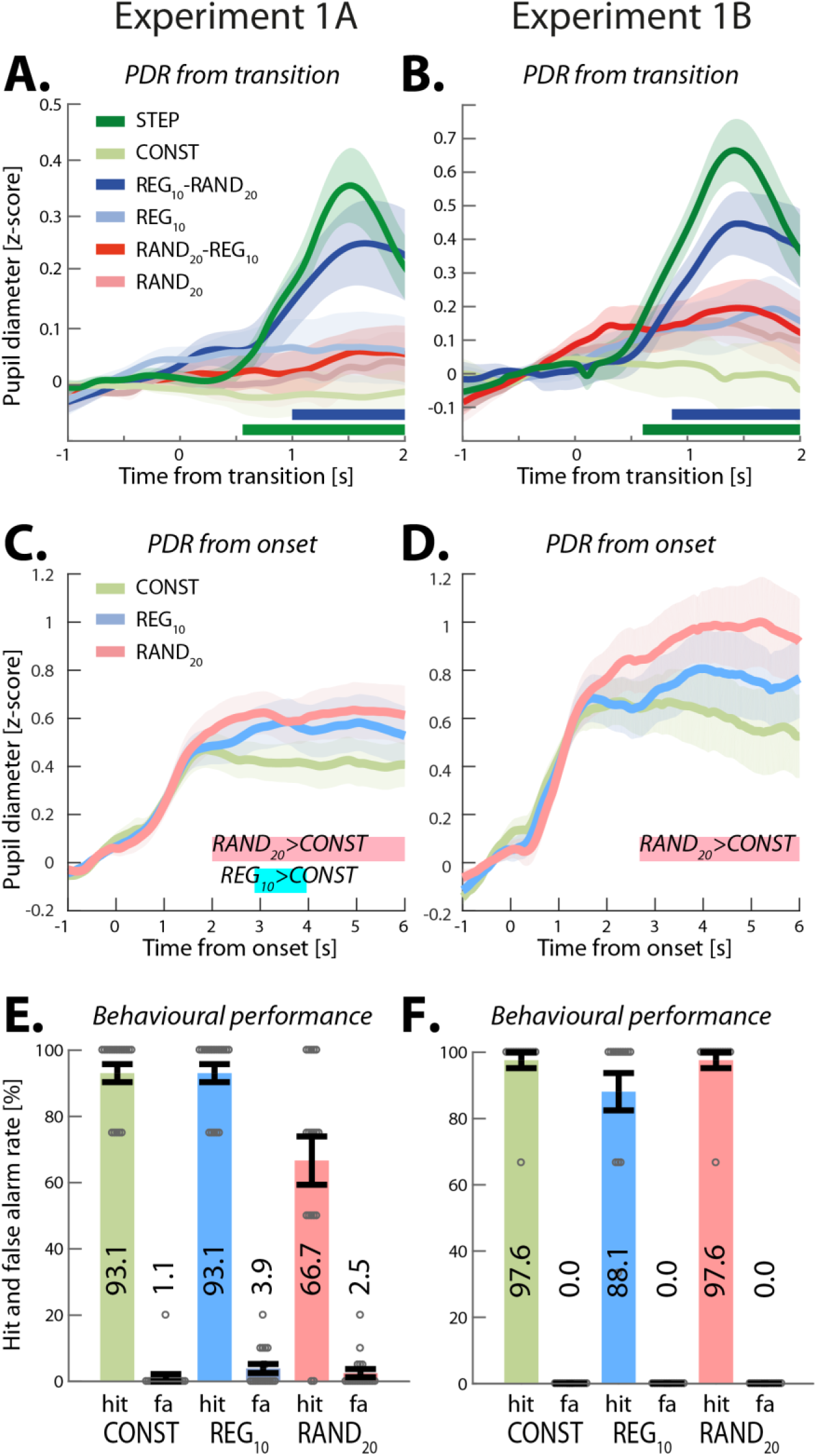
Experiment 1 (1A: N=18, 1B: N=14). REG-RAND but not RAND-REG transitions are associated with a pupil dilation response. [A] Average pupil diameter over time relative to the transition in Experiment 1A. Solid lines represent the average normalized pupil diameter. The shaded area shows ±1 SEM. Color-coded horizontal lines at graph bottom indicate time intervals where cluster-level statistics show significant differences between each change condition and its no-change control (p<0.05). In STEP, the pupil diameter started to increase around 300ms post-transition, reaching peak amplitude at 1520ms; it statistically diverged from its control, CONST, from 560ms through to sequence offset. Similarly, the PDR to REG_10_-RAND_20_ increased from ∼700ms post-transition, peaking at 1640ms. REG_10_-RAND_20_ statistically diverged from its control, REG_10_, at 1000ms post-transition and remained significantly higher until 2400ms. No significant differences between RAND_20_-REG_10_ and RAND_20_ were observed. Transition conditions were also compared directly (not shown): REG_10_-RAND_20_ was significantly higher than RAND_20_-REG_10_ from 840ms post-transition and up to stimulus offset, but no difference was observed between STEP and REG_10_-RAND_20_. [B] Average pupil diameter over time relative to the transition in Experiment 1B (replicating Experiment 1A). The divergence of STEP from its control, CONST, was significant from 600ms post-transition. REG_10_-RAND_20_ significantly diverged from its control, REG_10_, from 860ms. As in Experiment 1A, no significant differences were observed between RAND_20_-REG_10_ and RAND_20_ throughout the epoch. The difference between REG_10_-RAND_20_ and RAND_20_-REG_10_ began around 1020ms post-transition. REG_10_-RAND_20_ also showed a significantly greater PDR than STEP from 740ms onwards. [C, D] Average pupil diameter over time, relative to sequence onset. There were no significant differences between RAND_20_ and REG_10_ in either experiment. In Experiment 1B REG_10_ showed a significantly larger pupil diameter than CONST between 2880 and 3960ms post-onset. In both experiments, RAND_20_ was associated with a significantly larger pupil diameter than CONST from 2000ms (Expt1A) and from 2680ms (Expt1B) post-onset. Importantly there were no differences between RAND_20_ and REG_10_ in either experiment. [E] Behavioral hit and false alarm (fa) rates for the gap detection task in Experiment 1A. Grey circles represent individual participant data, and error bars are ±1 SEM. Performance on RAND_20_ was significantly reduced relative to the other conditions. [F] Gap detection task in Experiment 1B. Lengthening the gap in Experiment 1B resulted in equated performance across conditions.

Performance on the gap detection task was good overall (Fig. 2E) but we observed a main effect of condition on hit rates (arc-sine transformed for these and all subsequent statistical analyses on hit rates, F(1.198,20.372)=8.285, p=0.007). *Post hoc* tests confirmed that hit rate in RAND_20_ was lower than in CONST (p=0.026) and REG_10_ (p=0.026), while CONST and REG_10_ did not differ significantly (p=1.000). To assure that performance disparities were not driving differential PDR effects, gap duration in Exp. 1B was lengthened by 50ms to equate task performance across conditions.

### Exp. 1B (N=14)

The revised paradigm successfully eliminated performance differences between conditions (Fig. 2F), with a repeated-measures ANOVA showing no effect of stimulus condition on hits (F(2,26)=2.115, p=0.141) or false positive rates (F(2,26)=1.000, p=1.000).

The PDR pattern observed in Exp. 1A was entirely replicated (Fig. 2B). Overall, the results of Exp. 1 confirmed that PDRs are consistently evoked by STEP and REG_10_-RAND_20_ transitions, but not RAND_20_-REG_10_ transitions.

### The RAND-REG null effect is not due to pupillary saturation

It is important to eliminate the possibility that the lack of PDR to RAND_20_-REG_10_ transitions resulted from pupil diameter increase to saturation during the pre-transition RAND_20_ part of the sequence. Fig. 2C and D present average pupil diameter in the no-transition stimuli from sequence onset as measured in Exp. 1A and 1B. No significant differences were observed between RAND_20_ and REG_10_ in either Exp. 1A or 1B, suggesting an equivalent average pupil dimeter before the transition. Identical results were also obtained in Exp. 2 and 4 below (see Fig. 4E, 7B), confirming that the absence of a PDR in RAND_20_-REG_10_ transitions cannot be explained based on pre-transition differences between conditions. In Exp. 1 only there appears to be a pre-transition difference between either of the structured (RAND and REG) sequence conditions vs. the CONST condition. It may be due to the fact that the gap detection task was more demanding in the structured relative to the simpler sequences. However, this effect appears to be unstable (not observed with an identical task and stimuli in subsequent experiments see Fig. 4E, 7B) and is therefore not discussed further.

### The RAND-REG null-effect is not due to temporal spread of pupil dilation events

To confirm that the null effect for RAND_20_-REG_10_ indicates the absence of a pupil response and is not instead a consequence of an increased temporal spread of dilation events, pupil dilation (PD) and constriction (PC) rates were also analyzed (see Methods). This analysis is fundamentally different from the PDR analysis in that it focuses on the incidence of PD (or PC) events, irrespective of their amplitude, and therefore provides a sensitive measure of subtle changes in pupil dynamics potentially evoked by the transitions. Fig. 3 shows pupil dilation events from each trial, for each subject (N=32, combining Exp 1A & B), over an interval of 2 seconds before to 2 seconds after the transition.

**Figure 3:**
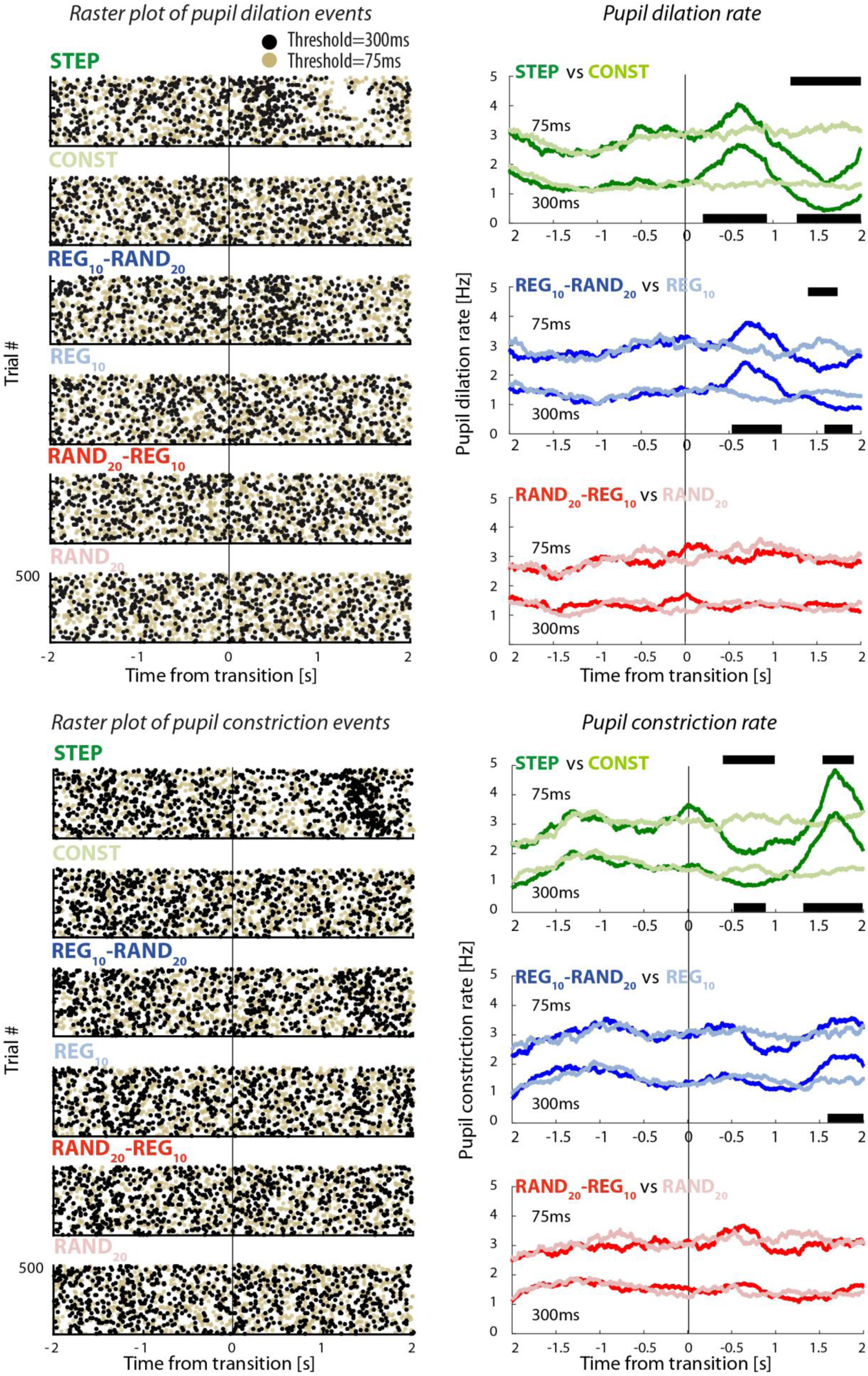
Experiment 1: Pupil dilation and constriction rates. [Top left] Raster plots of pupil dilation (PD) events extracted from all trials and all participants (collapsed over Experiment 1A & B). Each line represents a single trial. Black dots represent the onset of a pupil dilation with a duration of at least 300ms, yellow dots represent pupil dilation onsets with a threshold duration of 75ms. Transition time is indicated by a black vertical line. [Top right] Pupil dilation rate (running average with a 500ms window) as a function of time relative to the transition. The black horizontal lines indicate time intervals where cluster-level statistics showed significant differences between each change condition and its no-change control. The statistics for the PD rates with a threshold duration of 75ms and 300ms are placed above and below the graph, respectively. [Top] STEP and CONST; [Middle] REG_10_-RAND_20_ and REG_10_; [Bottom] RAND_20_-REG_10_ and RAND_20_. The lower panels present the pupil constriction (PC) rate results, arranged in the same format.

STEP and REG_10_-RAND_20_ transitions were associated with an increase in PD rate shortly after the transition, whereas no such change in rate was observed for RAND_20_-REG_10_. This effect was also mirrored in the constriction data, confirming that neither PD nor PC dynamics changed following RAND_20_-REG_10_ transitions. Results were equivalent across event-duration thresholds of 75 and 300ms (see Methods). Overall, this set of analyses provides further evidence for a null PDR response related to the RAND_20_-REG_10_ transition.

To interpret this first set of results, it is important to establish whether the PDR observed for STEP and REG_10_-RAND_20_ transitions revealed ‘true’ sensitivity to pattern violations, or rather was driven by low-level stimulus changes (frequency deviants). In Exp. 1, at least half of REG_10_-RAND_20_ trials involved the appearance of a novel frequency at the time of transition. This is also trivially the case for all STEP trials. It is therefore possible that the PDR reflects a simple response to the detection of a new frequency in the stimulus. In Exp. 2 the stimulus set was amended to include conditions where the transition was manifested as a change in pattern with or without frequency deviants.

## Exp. 2 (N=18): The pupil-linked LC-NE system is activated by ‘pure’ pattern violations

The stimulus set for Exp. 2 is in Fig. 4, top panel. Fig. 4B plots all the conditions which contained a regular-to-random transition: All evoked a marked PDR relative to the REG_10_ control. Notably a prominent PDR was observed for REG_10_-RAND_10_, i.e. a transition from a REG to a RAND pattern manifested as a change in pattern only, while maintaining the same 10 frequencies. In contrast, no significant difference was observed for any of the random-to-regular transitions (Fig. 4D). This was also the case for the RAND_10_-REG_10d_ condition where the RAND and REG sequences differed in frequency content in addition to the change in pattern. Whilst a small peak is visible in that condition, no significant differences are observed when compared to the no-transition RAND_10_ condition (Fig. 4C).

**Figure 4:**
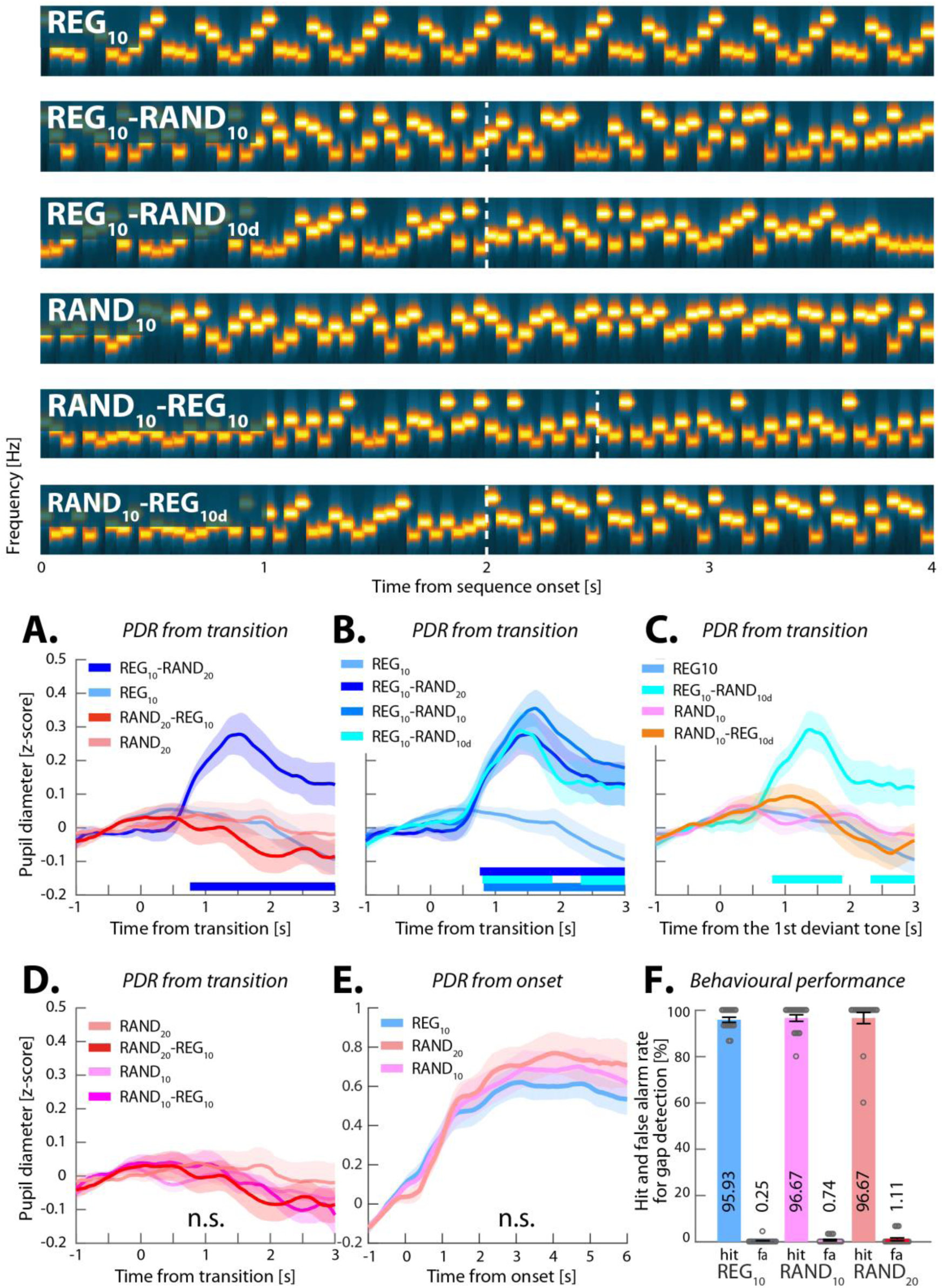
Experiment 2 (N=18). The PDR response reflects sensitivity to ‘pure’ pattern violations. [Top] Example spectrograms for the additional stimuli used in Experiment 2. REG_10_ and RAND_10_ were generated by randomly selecting 10 frequencies from the pool and then either iterating that sequence to create a regularly repeating pattern or presenting them in random order. REG_10_-RAND_10_ and RAND_10_-REG_10_ sequences were created from the same 10 frequencies (different sets on each trial). Thus, the transition was manifested as a change in pattern only, without the occurrence of a frequency deviant. REG_10_-RAND_10d_ and RAND_10_-REG_10d_ were created such that the frequencies used for the REG and RAND portions of the sequence were different (non-overlapping sets of 10 frequencies each). The transition was thus manifested as both a change in pattern, and also as a change in frequency content. The stimulus set also included REG_10_-RAND_20_, RAND_20_-REG_10_ and RAND_20_ sequences (identical to those in Experiment 1). Dashed vertical white lines indicate the transition time. Note that for RAND_10_-REG_10_ the transition time is defined as occurring after the first full regularity cycle (see also Figure 1). The transition time is not adjusted for RAND_10_-REG_10d_ because the transition becomes statistically detectable immediately when the alphabet changes (at the nominal transition time). For presentation purposes, the plotted sequence lengths are equal, but experimental sequences durations varied randomly between 6.0-7.5 s. [Bottom] [A-D] Average pupil diameter over time relative to the transition. Colored horizontal lines indicate time intervals where cluster-level statistics showed significant differences between each change condition and its no-change control. [A] The pupil responses to REG_10_-RAND_20_, RAND_20_-REG_10_ and their respective controls replicated the pattern observed in Experiment 1: REG_10_-RAND_20_ diverged from REG_10_ at 780ms post-transition; RAND_20_-REG_10_ did not differ statistically from RAND_20_. [B] A significant PDR was observed for all conditions containing transitions from REG_10_. The PDR in REG_10_-RAND_10_ increased from 420ms, and statistically diverged from REG_10_ 860ms post-transition, peaking at ∼1620ms; the PDR to REG_10_-RAND_10d_ diverged from REG_10_ at 800ms, peaking at 1380ms. [C] A comparison between the two conditions which contained a change in alphabet at the transition (REG_10_-RAND_10d_, RAND_10_-REG_10d_). Only REG_10_-RAND_10d_ evoked a PDR [D] None of the stimuli containing transitions from RAND_10_ evoked a significant PDR. [E] Average pupil diameter over time, from stimulus onset. No differences were observed between any of the conditions. [F] Behavioral results for the gap detection task. Error bars are ±1 SEM; grey circles represent individual participant data. Performance was at ceiling.

To confirm that the various REG and RAND conditions did not diverge pre-transition, we analyzed the pupil response from stimulus onset (Fig. 4E) and found no significant differences between any of the conditions. Consistent with Exp. 1, behavioral performance did not differ across conditions (Fig. 4F).

## Experiment 3 (N=14): The effect of active transition detection

Exp. 1 and 2 measured responses to transitions when they were not behaviorally relevant. To understand the effect of task relevance on PDRs, we introduced an active behavioral transition-tracking task. The experimental conditions were as in Exp. 1, but participants were asked to detect pattern changes rather than silent gaps.

**Behavioral results** are summarized in Fig. 5A. Hit rate data demonstrated that all transition conditions were highly detectable by human listeners. Although false positive rates were all low, there was a main effect of condition (F(1.405,18.262)=15.272, p<0.001), where, consistent with previous work (Barascud et al., 2016), there was a small but significantly higher false positive rate for (no-transition) RAND_20_ compared with CONST and REG_10_, with the latter two not differing significantly (p=0.083; Bonferroni corrected). To avoid confounds due to false positive disparities, all false positive trials were excluded from pupil analysis.

**Figure 5:**
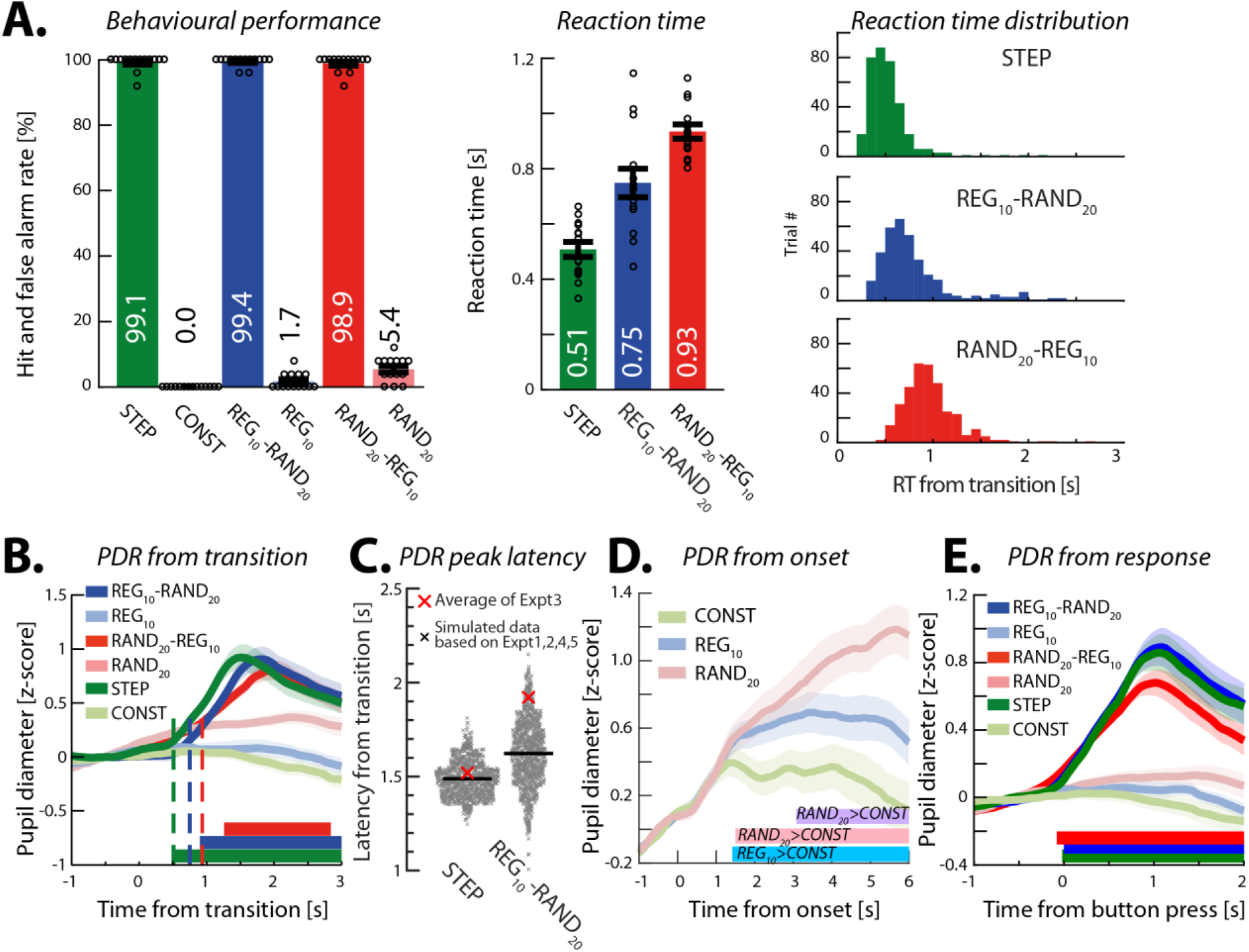
Experiment 3 (N=14): Active transition detection. [A] Behavioral performance (active transition detection). [Left] Hit rates and false positive rates. Circles indicate individual subject data; error bars are ±1 SEM. [Middle] Reaction times (RT). [Right] The distribution of RTs (across trials and participants) for each transition condition. The variance of the RT distribution of RAND_20_-REG_10_ was smaller than that of REG_10_-RAND_20_ (Levene’s test, F(1,690)=14.426, p=0.0002). [B] Average pupil diameter relative to the transition. Solid lines represent the average normalized pupil diameter, relative to the transition. Shading shows ±1 SEM. Colored horizontal lines indicate time intervals where cluster-level statistics showed significant differences between each change condition and its control. Dashed vertical lines mark the average RT for each condition. Clear PDRs were observed for all 3 transitions. The PDR to STEP increased from ∼490ms post-transition, peaking at 1500ms; it statistically diverged from CONST from 540ms through to sequence offset. For REG_10_-RAND_20_, the responses commenced ∼750ms post-transition, peaking at 1840ms, and statistically diverged from REG at 900ms post-transition through to sequence offset. For RAND_20_-REG_10_, the response started at 930ms, peaked at 1980ms, and statistically diverged from its control RAND_20_ between 1260 and 2840ms. Comparing transition conditions directly, no difference was observed between STEP and REG_10_-RAND_20_, but the pupil diameter in REG_10_-RAND_20_ became significantly greater than that in RAND_20_-REG_10_ between 700 and 1480ms post-transition. [C] A comparison of PDR peak latency to STEP and REG_10_-RAND_20_ in Experiment 3 relative to Experiments 1,2,4,5 (see methods). The scatterplots show the distribution of the simulated peak latency of STEP (left) and REG_10_-RAND_20_ (right) in the ‘gap detection’ experiments. The red crosses indicate the mean peak latency in Experiment 3. The results showed no difference (p = 0.338) for STEP, but a greater latency for REG_10_-RAND_20_ (p = 0.048). [D] Average pupil diameter over time relative to the sequence onset. RAND_20_ and REG_10_ statistically diverged from CONST at 1500 and 1420ms post-onset, respectively. Interestingly, RAND_20_ evoked an even larger PDR than REG_10_ from 3080ms post-onset. [E] Average pupil diameter over time, relative to button press.

For reaction time, a repeated measures ANOVA confirmed a main effect of condition (F(2,26)=90.723, p<0.001; STEP<REG-RAND<RAND-REG), consistent with previous work (Barascud et al., 2016).

Turning to **Pupillometry**, we observed three fundamental differences relative to Exp 1:

### (1) RAND-REG also evoked a PDR response

Fig. 5B plots the average normalized pupil diameter relative to the transition. Clear PDRs were observed in all three change conditions. Critically - and unlike the previous experiments with a gap detection task - a robust PDR was associated with RAND_20_-REG_10_ when the transition was task-relevant.

### (2) Delayed PDR peak for REG_10_-RAND_20_

Compared to Exp. 1 and 2, we also observed a substantial shift in the latency of the PDR to REG_10_-RAND_20_ (Fig. 5C). Active transition detection slowed the PDR by ∼300ms, with a peak latency of 1840ms in Exp. 3 relative to 1400-1600ms in the previous experiments. Peak latency to STEP did not change.

### (3) Differences in pupil diameter observed from sequence onset

In Exp. 1 and 2 (see also Exp. 4, below) we consistently observed no difference between REG_10_ and RAND_20_ when analyzing pupil responses relative to sound onset. In contrast, here in Exp 3 we observed a pre-transition disparity between REG_10_, RAND_20_ and CONST (Fig. 5D), such that RAND_20_ sequences evoked the largest sustained pupil dilation, followed by REG_10_. The sustained pupil diameter for RAND_20_ was significantly higher than REG_10_ from 3080ms post-onset. This cannot be explained by the higher false alarm rate of RAND_20_, as trials with incorrect responses were excluded from analysis, but may be a consequence of the computational demands or perceptual effort associated with tracking RAND_20_ sequences (see discussion).

We explored **the relationship between reaction time (RT) and pupil diameter across single trials** (Fig. 6). Generally, peak pupil dilation (indicated by hot colors in Fig. 6A) occurred about 1 second after the button press (Einhäuser et al., 2010). This relationship is also evident in Fig. 5E: the onset of the PDR to each of the transitions closely coincides with button press timing.

**Figure 6:**
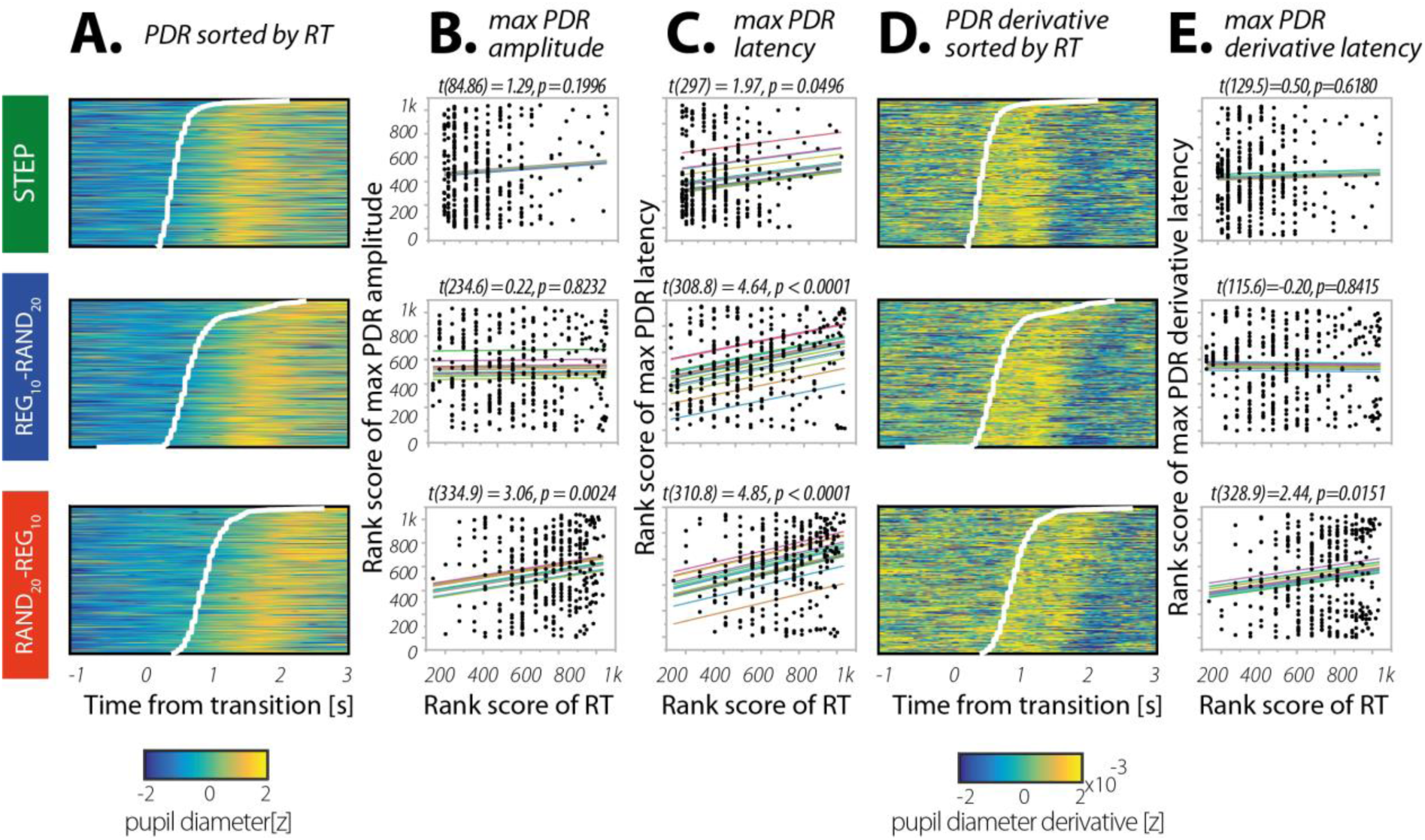
Relationship between RT and the pupil dilation response. [A] Single trials sorted by RT (y-axis, RT indicated by white lines) shown against the time relative to the transition (x-axis) with the colors showing pupil diameter (the warmer the color, the larger the pupil). [B] Scatter plots show the maximum PDR amplitude for each trial (ranked low to high on the y-axis) versus ranked RT for the same trial (x-axis), separated by condition as in [A]. Each line of fit shows the modelled random effect of subject (offset), with slope the fixed effect of RT. Fixed-effects t-values and associated p-values appear above each fitted scatterplot. [C] Rank maximum PDR latency vs. RT. Scatterplot and fitting as in [B]. [D] Single trials sorted by RT (y-axis, RT indicated by white lines) shown against the time relative to the transition (x-axis) with the colors showing the rate of change of pupil diameter (the warmer the color, the larger the rate of change in pupil size). [E] Rank maximum derivative latency vs rank RT (scatterplot fits as in [C]).

The fact that peak PDR occurs substantially after the button press may indicate that the behavioral response itself either triggers or otherwise modulates the PDR. To understand the relationship between RT and the PDR, we examined whether button press timing was systematically linked to key measures of PDR dynamics. We tested the **relationship between RT and the maximum PDR amplitude** (Fig. 6B) on a single-trial basis. (All subsequent analyses are rank-based, with subjects as random effect and reaction time as fixed effect (Bates et al., 2014, see Methods)). In the STEP condition, there was no significant association between RT and maximum PDR amplitude (t(84.86) = 1.29, p = 0.1996); this also held true in the REG_10_-RAND_20_ condition (t(234.6) = 0.22, p = 0.8232). However, in the RAND_20_-REG_10_ condition, RT was significantly associated with maximum PDR amplitude (t(334.9) = 3.06, p = 0.0024): here, RT was estimated to account for 2.8% of variance in the maximum PDR amplitude (estimate of partial R^2^ derived using an implementation of the Nakagawa and Schielzeth (2013) algorithm, see Methods).

We then asked **whether RT was associated with the timing of the maximum PDR** (Fig. 6C). In the STEP condition, there was a small but significant association between RT and PDR latency (t(297) = 1.97, p = 0.0496, accounting for an estimated 1.5% of PDR latency variance. (Note, however, this association was not significant when slopes were allowed to vary). In the REG_10_-RAND_20_ condition, RT was also significantly associated with PDR latency (t(308.8) = 4.64, p < 0.0001), and accounted for an estimated 7.2% of its variance. Finally, RT was also associated with PDR latency in the RAND_20_-REG_10_ condition (t(310.8) = 4.85, p < 0.0001), and an estimated 6.3% of RT variance.

Finally, we asked **if RT was associated with the timing of the maximum derivative of the PDR** (i.e. the time at which the rate of change of pupil size is maximal; Fig. 6E). As with the other two dependent variables, reaction times in STEP were not associated with the maximum derivative PDR latency, t(129.5)=0.50, p=0.6180). Nor was there a significant association in the REG_10_-RAND_20_ condition (t(115.6)=-0.20, p=0.8415)). However, RT in the RAND_20_-REG_10_ condition was associated with the maximum derivative PDR latency, t(328.9)=2.44, p=0.0151), and 1.8% of its variance.

In sum, the pattern of associations between participants’ reaction times and various pupil measures suggests that the amplitude and timing of the pupillary response to both RAND_20_-REG_10_ and REG_10_-RAND_20_ are related to button press timing, but only to a modest degree, with RT accounting for between ∼2-6% of estimated variance. While these analyses are limited by the relatively small amount of trials per condition/subject, this outcome suggests that the appearance of a PDR to the RAND_20_-REG_10_ transition in Exp 3 is not primarily driven by the button press. Rather, having listeners actively monitor and respond to the statistical transitions prompted a change in the underlying cognitive process, e.g. by delineating the category (or decision) boundary between RAND and REG, and thereby rendering the transition as a model violation (see Discussion).

## Exp 4: Pupil responses to transitions from randomness

We have argued that to achieve effective model maintenance, the brain must arbitrate between gradual and punctate environmental changes. In other words, at each point in time, the brain must decide whether to continue updating its current representation of environmental contingencies or instead abandon the existing model and prioritize bottom-up evidence accumulation (“out with the old, in with the new”). Our results thus far suggest that what determines the difference between gradual and abrupt change can be gleaned through delineating the contingencies which evoke a PDR.

In REG-RAND, and trivially so in STEP, the statistical violation is immediately observable if listeners form a robust representation of the patterning of the REG sequences This could therefore be sufficient to trigger the abrupt-model violation signal.

By contrast, since absolutely any sequence of tones has the same probability under RAND, the detection of transitions out of this distribution is statistically more complicated. The lack of a PDR for RAND_20_-REG_10_ transitions, when not behaviourally relevant, is taken to indicate that that this transition is indeed not treated as an abrupt model violation. In Exp 4 we explore whether the same is true for less complex regular patterns.

### Exp 4A (N=12)

We asked whether the most basic regular pattern - a single repeating tone (REG_1_) - evokes a PDR. Naive listeners (performaing a gap detection task) were presented with RAND_20_-REG_1_ sequences (Fig. 7, top panel), in addition to STEP, REG_10_-RAND_20_ and RAND_20_-REG_10_. Replicating the result of Exp 1, we observed a PDR to REG_10_-RAND_20_ but not to RAND_20_-REG_10_. By contrast, the RAND_20_-REG_1_ transition evoked a fast-onset and robust PDR of similar amplitude to that evoked by transitions from regularity to randomness. This result is consistent with previous demonstrations that the violation of randomness by repetition evokes an MMN-like response (Horváth and Winkler, 2004; Rosburg, 2004) - a finding which was taken to suggest that the auditory system represents stochastic frequency variation as a regularity *per se* (Wolff and Schröger, 2001).

**Figure 7:**
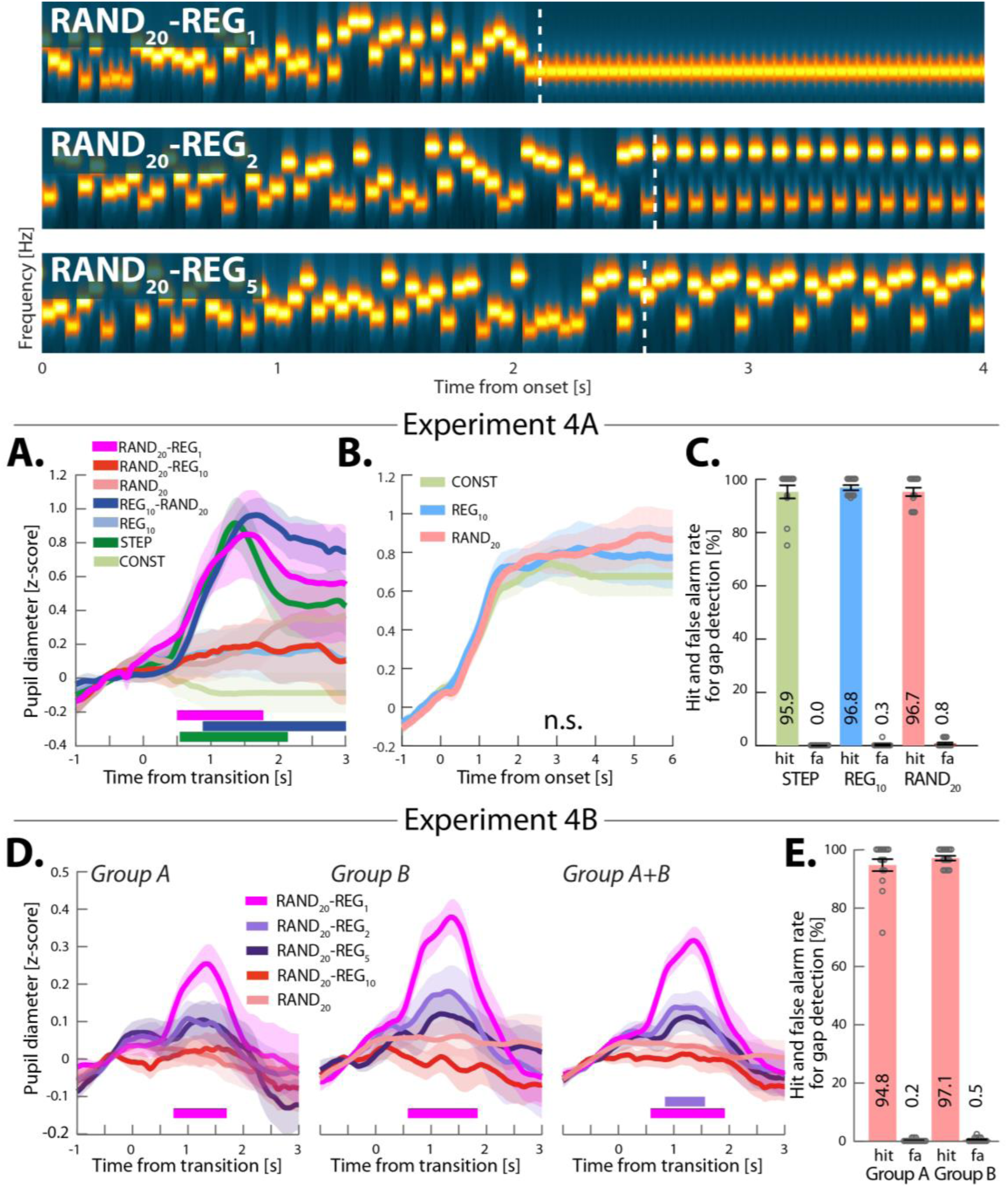
Experiments 4A (N=12) and 4B (two groups of N=15). Abrupt reduction in the PDR for regularities more complex than REG1. [Top] Example spectrograms for the additional stimuli used in Experiments 4A and B. RAND_20_-REG_1_ consisted of a transition from a random sequence (generated by sampling frequencies from the full pool with replacement RAND_20_) to a single repeating tone. RAND_20_-REG_2_ consisted of a transition from a RAND_20_ sequence to a regular pattern consisting of two randomly selected tones. RAND_20_-REG_5_, consisted of a transition from a RAND_20_ sequence to a regular repeating pattern consisting of 5 tones. Dashed vertical white lines indicate the transition time, defined as occurring after the first full regularity cycle. For presentation purposes the plotted sequence lengths are equal. Actual durations varied randomly between 6 to 7.5s. In Experiment 4A the stimulus set also contained RAND_20_-REG_10_, REG_10_-RAND_20_, STEP, REG_10_, RAND_20_ and CONST sequences. In Experiment 4B the stimulus set additionally contained RAND_20_-REG_10_ and RAND_20_ sequences. [Bottom] [A] Average pupil diameter relative to the transition in Experiment 4A. Solid lines represent the average normalized pupil diameter, relative to the transition. The shaded area shows ±1 SEM. Colored horizontal lines indicate time intervals where cluster-level statistics showed significant differences between each change condition and its control. A robust PDR was evoked by the transition in RAND_20_-REG_1_, becoming significant from 500ms post-transition, at approximately the same time as STEP, and lasting through to 1780ms. The data also replicate the general pattern in Experiments 1 and 2: a PDR evoked by REG_10_-RAND_20_ from 860ms onwards, and no significant difference between RAND_20_-REG_10_ and RAND_20_. The transition conditions were also compared directly and revealed a significant difference between REG_10_-RAND_20_ and RAND_20_-REG_10_ from ∼1020ms post-transition. [B] Average pupil diameter over time from stimulus onset. No differences were observed between any of the conditions. [C] Behavioral results for the gap detection task in Experiment 4A with ±1 SEM error bars, and grey circles representing individual participant data. There was no statistical difference between conditions. [D] Average pupil diameter over time relative to the transition in Experiment 4B: [Left] Group A (N=15). A clear PDR is observed for RAND_20_-REG_1_ which diverged from its control, RAND_20_, from 740 to 1700ms. No significant differences were observed in the other conditions. [Middle] Group B (N=15). RAND_20_-REG_1_ diverged from RAND_20_ between 580ms-1840ms; RAND_20_-REG_2_, RAND_20_-REG_5_ and RAND_20_-REG_10_ were not significantly different from RAND_20_. [Right] both groups combined (N=30). Significant PDRs were observed for RAND_20_-REG_1_ (from 580 to 1920ms) and RAND_20_-REG_2_ (from 840 to 1560ms) transitions only. [E] The behavioral performance for both groups was at ceiling.

As with the previous experiments, there was no significant difference between REG_10_ and RAND_20_ pre-transition (Fig. 7B), and no behavioral difference in the gap detection task across the three conditions (Fig. 7C).

### Exp 4B (two groups of N=15)

How does the PDR evolve as the regularity becomes more complex, e.g., as we add more elements to the regular pattern? In Exp 4B, in addition to RAND_20_-REG_1_, we also included transitions from RAND_20_ to repeating 2-, 5-, and 10-tone patterns (REG_2_, REG_5_, REG_10_, Fig. 7 top panel, REG_10_ not shown).

Due to the increased number of conditions and therefore longer experiment time, data were noisier than in the previous experiments. Thus, to replicate the effects from the first group (group A; N=15), the experiment was repeated in another group of listeners (group B; N=15). We also collapsed the data across both groups to maximize statistical power.

For both groups, the behavioral performance in the gap detection task was at ceiling (Fig. 7E). Fig. 7D plots the results of Exp 4B for each group separately, and when pooled together. In each group, and as in Exp 4A, RAND_20_-REG_1_ evoked a significant PDR; however, a clear PDR was not observed for the other conditions. The average normalized pupil diameter in RAND_20_-REG_2_ and RAND_20_-REG_5_ appeared to suggest a small (non-significant) gradual increase between 640ms and 800ms, which might indicate the presence of a sub-threshold effect. Indeed, collapsing the data across groups did demonstrate a significant PDR for RAND_20_-REG_2_ between 840–1560ms post-transition. The PDR for RAND_20_-REG_5_ remained non-significant. These effects are also mirrored in the PD rate analysis (see sup. Materials).

Overall the results of Exp 4B demonstrate a sharply reduced PDR for regularities more complex than REG_1_: While REG_1_ consistently evoked a robust response in both groups A and B, and with an amplitude identical to that observed for STEP and REG_10_-RAND_20_ (Exp 4A, Fig. 7A), the PDR to RAND_20_-REG_2_ was substantially reduced, such that it required double the power to reach significance. The small effect of REG_2_ may indicate that for some subjects or in a subset of trials a PDR was present. The lack of a PDR for more complex regularities suggests that the associated transitions are treated as a gradual rather than abrupt transition with respect to the internal model maintained for RAND_20_.

## General Discussion

We report two main findings: First, when pattern changes were behaviorally irrelevant, pupil dilation responses (PDR) were evoked exclusively by changes associated with violations of regularity. Second, behavioral relevance exerted a major effect on pupil dynamics, changing the responses both during the establishment of patterns, and at transition periods.

Research in animal models has established a robust link between phasic pupil responses and spiking activity within the LC, providing compelling evidence for pupil dynamics as an indirect measure of NE release (Joshi et al., 2016). Our observations are therefore interpreted in the context of understanding the role of the pupil-linked LC-NE system in reporting on aspects of the statistics of rapid sensory signals. The results are consistent with a hypothesized role of NE as a model interrupt signal, and provide a rich view of the contingencies that have automatic and/or controlled access to this interrupt.

A large body of work has suggested a gating role for NE in balancing bottom-up-driven sensory processing vs. top-down priors (Bouret and Sara, 2005; Dayan and Yu, 2006; Sara and Bouret, 2012; Yu and Dayan, 2005). Direct electrophysiological recording in animal models (Aston-Jones et al., 1997; Bouret and Sara, 2004; Sara and Segal, 1991; Vankov et al., 1995) and fMRI in humans (Payzan-LeNestour et al., 2013) have observed neural activity in LC in response to unexpected and abrupt contextual changes. Pharmaceutical evidence has demonstrated that downregulating NE results in impaired adaptation to environmental changes (Marshall et al., 2016; McGaughy et al., 2008) whereas pharmacologically stimulating the noradrenergic system is associated with increased learning rates (Devauges and Sara, 1990; Howlett et al., 2017). Indirect measures of NE release, based on pupillometry, have also revealed an association between NE signaling and increased learning rates – a proxy for model resetting (Krishnamurthy et al., 2017; Nassar et al., 2012). However, the existing literature is limited by the fact that most of the experimental results taken to support the ‘NE as an interrupt signal’ hypothesis have involved tasks in which inputs evolve slowly over time and participants make either active decisions about stimulus predictability or are required to form stimulus – response associations (Jepma and Nieuwenhuis, 2011; Krishnamurthy et al., 2017; Lavín et al., 2014; Marshall et al., 2016; Nassar et al., 2012; Payzan-LeNestour et al., 2013; Preuschoff et al., 2011). Here we demonstrate that NE also plays a role in coding model violations on time scales relevant to tracking unfolding sensory information, even when it is not behaviorally relevant.

### PDR specificity under behaviorally-irrelevant listening conditions

Deviants or salient changes in sound sequences are well known to evoke PDRs, even under passive listening conditions (Liao et al., 2016; Nieuwenhuis et al., 2011). These observations have prompted a suggestion that, as part of a broader ‘fight or flight’ response, pupil activation reflects the operation of an interrupt signal that halts current ongoing activities to allow an attentional shift towards the new event, thus facilitating adaptive behavior (Nieuwenhuis et al., 2011; Wang and Munoz, 2015). Phasic LC-NE activation has duly been hypothesized to serve as a neural interrupt signal for unexpected events (Bouret and Sara, 2005; Dayan and Yu, 2006; Payzan-LeNestour et al., 2013), prompting the resetting of top-down connectivity when sensory information indicates that the currently instantiated model of the environment is no longer valid. These observations raise the obvious - and behaviorally important - question of what exact changes are able to drive the interrupt signal.

One common idea is that sensory processing involves rich statistical modeling (Friston, 2005; Knill and Richards, 1996; Rao, 2005), which suggests that many changes might drive the interrupt signal. However, not only are there obvious dangers to making interruption too promiscuous when neural processing is focusing on a task for which the statistics are irrelevant, but it is also computationally costly to build detailed models of complex signals when these do not matter. This consideration motivated a heuristic separation between expected and unexpected uncertainty (Dayan and Yu, 2006; Yu and Dayan, 2005) with events falling in the latter category triggering model interruption, allowing a more sophisticated model-building process to occur, if appropriate.

Here, we exploited the statistical asymmetry between REG-RAND and RAND-REG to gain some further insight into these limits. The transition in REG-RAND is simple to detect, given knowledge of REG. However, for the RAND-REG transition, the RAND model is not directly falsified because all tones within the REG pattern are strictly consistent with the RAND model. The hypothesis is therefore that they are treated in terms of expected - rather than unexpected - uncertainty, leading to gradual rather than abrupt model change. In line with the formulation proposed by Dayan & Yu (2006), we found that, when the transition was irrelevant to the behavioral goal, the LC-NE system appeared to ignore RAND-REG transitions. This was observed for even moderately complex REG patterns, and even though listeners - and their brain activity - readily detect them on a similar time scale to REG-RAND transitions. This not only refutes the suggestion that any perceptually salient and contextually novel set of observations can drive NE, but also hints at limits to the statistical model-building process.

It is important to note that our stimuli were matched spectrally, and in terms of behavioral detectability and time to process, and were presented at a rate beyond that at which listeners can actively track structural content (Warren, 2008). Thus we extend to the preattentive case findings about NE and interrupts originally derived from decision-making tasks and demonstrate that the pupil-linked LC-NE system plays an obligatory role in tracking the statistics of unfolding sensory input.

The sharp drop in the PDR in RAND_20_-REG_2_ relative to RAND_20_-REG_1_ (Fig. 7D), indicates that, while the transition in RAND_20_-REG_1_ is coded explicitly as an abrupt model violation, the transition in RAND_20_-REG_2_ - from a random to a two-tone pattern - is not. One suggestion is that the brain engages in a form of automatic latent model building using just the last few tones. If the latent model based on those few tones fits them much better than the prevailing model, then an abrupt change is reported. Under this hypothesis, the fact that even as simple a sequence as two alternating tones does not generally lead to model change detection suggests stringent constraints on the automatic model construction – perhaps that it encompasses no more than two successive tones. It is tempting to speculate that such model construction could be implemented by low-level coding mechanisms e.g. adaptation or repetition suppression, both of which would lead to detectably unusual patterns of activity in tonotopically organized neural populations.

Importantly, we interpret the abrupt difference between RAND-REG_1_ and RAND-REG_2_ (despite their perceptual and statistical similarity) as further evidence for the specificity of the PDR for punctate changes in stimulus statistics. Critically, this specificity can be reversed when transitions are behaviorally relevant (Exp. 3; see more below).

### Relationship to brain responses

The observed PDR specificity is consistent with patterns of brain responses measured with Electro- and Magneto-encephalography (EEG, MEG) in naïve, distracted listeners (Fig. 1B). Robust brain responses are observed to both RAND-REG and REG-RAND transitions, but importantly, the response dynamics are distinct, revealing the differing computational demands of each transition type: RAND-REG transitions evoke a progressive increase in sustained brain responses, hypothesized to reflect the gradual increase in model precision associated with the increased predictability of the REG patterns (Barascud et al., 2016). This rise in the sustained response is underpinned by a distributed brain network of auditory cortical, frontal and hippocampal sources (Barascud et al., 2016; Southwell et al., 2017). Together, these sources are hypothesized to reflect the instantiation of a top-down model, producing increasingly reliable priors for upcoming sounds (Auksztulewicz et al., 2017).

In contrast, REG-RAND transitions evoke a mismatch response (similar in its dynamics to the MMN; Näätänen et al., 2007), followed by an abrupt drop in sustained activity, in line with immediate suppression of top-down prior expectations (see Barascud et al, 2016). The activity then settles at a low sustained level, consistent with the far weaker statistical constraints in the RAND pattern.

The PDR results point to a potential role for NE signaling in supporting the MEG-indexed ‘resetting response’ observed during REG-RAND transitions. The relevant neural circuit may involve signaling from MMN-related brain systems (Auditory Cortex and right IFG; Garrido et al., 2008) to the LC-NE system, possibly via the ACC (Behrens et al., 2007; Karlsson et al., 2012; Payzan-LeNestour et al., 2013) or orbitofrontal cortex (Nogueira et al., 2017; Southwell and Chait, 2018). LC activation would then trigger NE-mediated rapid interruption of the temporo-frontal network associated with generating top-down prior expectations. Further investigation combining pupillometry and sensitive source imaging are necessary to identify these circuits, and to test the proposed linked between the MMN response and NE release.

Notably, the timing of the effects observed here implicate rapid signaling from the auditory system to the LC. The onset of the PDR observed here is at ∼500ms post transition; PDRs evoked by simple salient auditory or visual stimuli are commonly estimated to onset at ∼300ms (Hoeks and Levelt, 1993; Wang and Munoz, 2015), suggesting that the REG-RAND transition-related signal reaches the pupil no later than ∼200ms post transition. This latency is within the same range as the abrupt drop in MEG activity recorded from auditory cortex (Fig. 1B), thereby temporally aligning pupil dilation time with the putative signature of NE release.

### Substantial influence of attentional set on pupil responses to change in sequence structure

Rendering the sequence transitions behaviorally relevant resulted in marked differences in pupil response dynamics. Notably, active monitoring of transitions gave rise to a PDR to RAND_20_-REG_10_ transitions and introduced a large delay (300ms) to the PDR to the REG_10_-RAND_20_ transitions. These effects were not strongly linked to the execution of a motor command, as evidenced by the fact that RT accounted for relatively little variance in various PDR metrics. Therefore, these behavior-related changes in the PDR to RAND_20_-REG_10_ and REG_10_-RAND_20_ likely reflect a change in the functional state of the LC-NE system, or inputs to it. This may be a consequence of a behaviorally-driven emergence of a category boundary between REG and RAND, a richer representation of the statistics of the patterns before or after the transition, or a threshold change for model reset. That the STEP transition appears unaffected may be because this change can be detected relatively early in the auditory processing hierarchy, and thus may not depend on feedback information flow in the same way as do REG_10_-RAND_20_ and RAND_20_-REG_10_.

We also observed a change in the dynamics of tonic pupil activity, i.e. in response to the ongoing sequence before the transition. When RAND and REG states were behaviorally irrelevant (in all but the third experiment), we observed no difference between the ongoing response to REG and RAND throughout the entire epoch (Fig. 2C,2D,4E,7B). However, making transitions between these states behaviorally relevant resulted in diverging PDRs to the different conditions themselves, even in the absence of a transition (Fig. 5B and 5D). Notably, these differences were observed even though the statistical structure *per se* is not explicitly trackable by listeners, due to the rapid rate at which successive tones are presented. Previous work has linked tonic pupil diameter differences to representation of expected uncertainty (Krishnamurthy et al., 2017; Nassar et al., 2012) possibly driven by cholinergic signaling (Reimer et al., 2016; Yu and Dayan, 2005). The present effects may be consistent with this interpretation; as indeed RAND_20_ is associated with less reliable priors than REG_10_. However, the fact that these differences in pupil diameter were observed exclusively during the active change detection task must therefore suggest that cholinergic activation is dependent on behavioral relevance and is not involved in automatic tracking of sequence predictability. An alternative, but not mutually exclusive, possibility is that this effect may reflect heightened vigilance or listening effort (Zekveld et al., 2014) arising through active sequence structure scanning, which is more demanding for RAND_20_ (Southwell et al., 2017).

### Conclusions

The data reported here demonstrate that the pupil-linked LC-NE system plays an obligatory role in tracking and evaluating the statistics of unfolding sensory input, thereby supporting brain networks involved in maintaining flexible perceptual representations in changing environments. However, this system is confined in the circumstances under which it signals an interrupt, particularly in the absence of a remit from a task. Together with previous work in the decision-making and learning fields, the present results establish a unified view of NE as a model interrupt signal operating on multiple time scales, from those relevant to tracking reward environments in the context of decision making to tracking rapidly-unfolding sensory environments during perception.

## Acknowledgements

We are grateful to Alex Billig and Shihab Shamma for comments and discussion and to Makoto Yoneya (NTT) for initiating the pupillometry setup at UCL. This work was supported by a BBSRC project grant and a BBSRC international partnering award to MC; SZ was partly funded by an NTT-UCL enhanced research contract. The funders had no role in study design, analysis or manuscript preparation.

## Author Contributions

**SZ** conceived performed and analyzed the experiments; wrote manuscript.

**MC** supervised and administered the project; secured the funding; conceived the experiments; wrote the manuscript.

**FD** conceived the experiments; formal analysis of Experiment 3; wrote the manuscript.

**PD** provided expertise and feedback; wrote the manuscript.

**SF** provided expertise and feedback; commented on manuscript draft.

**HL** performed pilot experiments; provided expertise and feedback; commented on manuscript draft.

## Declaration of Interests

The authors declare no competing interests.

## Methods

### CONTACT FOR REAGENT AND RESOURCE SHARING

Further information and requests for resources and data should be directed to and will be fulfilled by the Lead Contact: Maria Chait

### SUBJECT DETAILS

#### Ethics declaration

The experimental procedures were approved by the Research Ethics Committee of University College London. Participants were provided written informed consent and were paid for their participation.

#### Participant exclusion criteria and justification of N

The following exclusionary criteria were consistently applied across all experiments: To ensure that observed changes in pupil diameter were not blink-related artifacts, participants were excluded if they blinked in more than 50% of trials. Additionally, participants were excluded if their mean gaze location exceeded three standard deviations from the group mean.

The experiments were not conducted in the order in which they are reported. Initial experiments involved larger numbers (N=20). Over time, and as the first author became more experienced with conducting the experiments, it was concluded that smaller N and fewer trials per condition were sufficient.

#### Participant details

All participants reported normal hearing, normal or corrected-to-normal vision, and no history of neurological disorders.

##### Experiment 1A

Data from 18 participants (11 females; aged 20–29, average 23.41) are presented. Data from one additional participant were excluded due to failure to complete the experiment. Two further participants were excluded due to high blink rates in the STEP condition.

##### Experiment 1B

Data from 14 new participants (13 females; aged 22-26, average 23.1) were used in the analysis. Five additional participants were excluded due to high blinks rates. One further participant was excluded due to poor behavioral performance (0% gap detection hit rate in REG sequences).

##### Experiment 2

Data from 18 new participants (15 females; aged 20-35, average 25.1) are reported. Two additional participants were excluded: one due to high blink rates, and one due to wandering gaze.

##### Experiment 3

Data from 14 participants (10 females; aged 22–30, average 24.3) are presented. None were excluded.

##### Experiment 4A

Data from 12 new participants (9 females; aged 21 - 26, average 23.6) are presented. None were excluded.

##### Experiment 4B

This experiment was performed twice; a total of 30 new participants took part, with 15 participants initially (11 females; aged 20–29, average 23.5) and a new group of 15 participants subsequently (14 females; aged 20–25, average 22.5) to replicate the results of the first cohort. None were excluded.

### METHOD DETAILS

#### Pupil size measurement and analysis

Participants sat in front of a monitor at a viewing distance of 60cm in a dimly lit, acoustically shielded room (IAC triple walled sound-attenuating booth), with their head supported on a chinrest. They were instructed to continuously fixate at a white cross presented at the center of the screen (BENQ XL2420T with resolution of 1920×1080; refresh rate of 60 Hz) against a black background. The visual display remained constant throughout the session. An infrared eye-tracking camera (Eyelink 1000 Desktop Mount, SR Research Ltd.), positioned just below the monitor, continuously tracked gaze position and recorded pupil diameters, focusing binocularly at a sampling rate of 1000 Hz. The standard five-point calibration procedure for the Eyelink system was conducted prior to each experimental block. Participants were instructed to blink naturally.

Only the left eye was analyzed. To avoid contamination by blinks, which tended to increase towards the end of the stimulus, the final 0.5 s of each trial were cut from the analysis. The epochs therefore spanned one second before to two seconds post transition (Experiment 1) or three seconds post transition (all other experiments). This cut-off was comfortably beyond the time needed to detect the transitions, as corroborated by behavioral and MEG results (Experiment 3 and Barascud et al., 2016). Data in each epoch were smoothed with a 150ms Hanning window, z-scored for each block and baseline-corrected by subtracting the median pupil size of the pre-transition baseline. Intervals where the eye tracker detected full or partial eye closure were automatically treated as missing data and recovered with shape-preserving piecewise cubic interpolation. Trials with more than 50% missing data were excluded from analysis (<2 trials per subject). The normalized pupil diameter was time-domain-averaged across all epochs of each condition type to produce a single time series for each condition. Matched no-transition conditions were epoched in a similar manner around ‘dummy’ transition times set to match those in the transition conditions. To compare pupil dynamics from sequence onset, the data in the no change conditions (REG, RAND, CONST) were epoched from 1 second before sequence onset to 6 seconds post-onset and processed as described above.

#### Pupil event rate analysis

Pupil event rate analysis compared the incidence of pupil dilation or constriction events. Following Joshi et al, (2016), events were defined as local minima (dilations; PD) or local maxima (constrictions; PC) with the constraint that continuous dilation or constriction is maintained for at least 75ms (yellow dots in Figure 3) or 300ms (black dots in Figure 3). Both thresholds provided consistent data (see results), as did intermediate thresholds (including 100ms and 125ms; not shown). The rate was estimated for each subject separately by using a sliding 500ms window over all trials in each condition and comparing rate changes across time and condition (see ‘Statistical Analysis’, below). This relatively long window enabled us to capture possible subtle changes in the rate of occurrence of PD events. The analysis interval was between 2 seconds before to two seconds after the transition. Previous work (Barascud et al., 2016) demonstrated that brain responses to the transitions occur within < 300ms and behavioral responses (button press) are completed by 1000ms (see also Exp. 3 below), thus suggesting that the analysis interval is appropriate for revealing any effects. We also estimated rate by tallying PD or PC events with non-overlapping 500ms windows, and by convolving with an impulse function (see also Joshi et al., 2016; Rolfs et al., 2008). For each condition, in each participant and trial, the event time series were summed and normalized by the number of trials and the sampling rate. Then, a causal smoothing kernel ω(τ) = *α*^2^ × τ × *e*^−*α*τ^ was applied with a decay parameter of 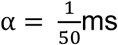
(Dayan and Abbott, 2001; Rolfs et al., 2008; Widmann et al., 2014). All analyses yielded identical results, therefore only the former is reported.

#### Experiment 1A Stimuli and procedure

The stimuli were sequences of concatenated tone-pips (50ms) with frequencies drawn from a pool of 20 fixed values (log-spaced) between 200 and 2000 Hz. The tone-pips were arranged according to six frequency patterns, generated anew for each participant (Figure 1): **CONST** sequences consisted of a single repeating tone, chosen by randomly selecting a frequency from the pool on each trial; **STEP** sequences consisted of a step change from one repeating tone to another repeating tone of a different frequency (both frequencies randomly drawn on each trial); **REG**_10_ sequences were generated by randomly selecting (with replacement) 10 frequencies from the pool and then iterating that sequence to create a regularly repeating pattern (with new patterns generated on each trial); **RAND**_20_ sequences were generated by randomly sampling frequencies from the pool with replacement; **REG**_10_**-RAND**_20_ and **RAND**_20_-**REG**_10_ sequences contained a transition between a regular pattern and a random pattern. The stimulus length varied randomly between 5 and 7 seconds, with a jittered transition time at around 2.5- and 3.5-seconds post-onset.

Sounds were presented diotically through headphones (Sennheiser HD558) via a Creative Sound Blaster X-Fi sound card (Creative Technology, Ltd.) at a comfortable listening level, self-adjusted by each participant. Stimulus presentation and response recording were controlled with the Psychtoolbox package (Psychophysics Toolbox Version 3; Brainard, 1997) in MATLAB (The MathWorks, Inc.). 146 stimuli – 24 for each condition - were randomly presented in four consecutive blocks (separated by 3 min breaks) with an inter-trial interval of six seconds. 25% of the signals contained a silent gap, occurring at any time from 250ms post onset to 750ms pre-offset. Participants were instructed to monitor the sequences for these events and to respond by pressing a button as quickly as possible. To equate for task difficulty, the gap consisted of one missing tone (50ms) in the CONST and STEP sequences, and two missing tones (100ms) in REG and RAND sequences. Visual feedback, lasting 400ms, was provided immediately at the end of each sequence. Trials containing a gap and trials in which participants made a false positive were excluded from further analysis.

#### Experiment 1B Stimuli and Procedure

The stimuli and procedure were identical to Experiment 1A, except that only three blocks were run, for a total of 108 stimuli. The data from Experiment 1A indicated that this was sufficient to measure the relevant effects. To address behavioral differences between conditions observed in Experiment 1A, the gap in RAND sequences was lengthened to 3 tones (150ms).

#### Experiment 2 Stimuli and Procedure

The stimulus set is described in Figure 4. Stimulus length was randomly varied between 6-8 seconds, with the transition jittered between 3-4 seconds after sequence onset. A total of 240 stimuli were presented in random order over 5 consecutive blocks: 60 REG_10_, 20 REG_10_-RAND_10_, 20 REG_10_-RAND_10d_, 20 REG_10_-RAND_20_, 40 RAND_10_, 20 RAND_10_-REG_10_, 20 RAND_10_-REG_10d_, 20 RAND_20_ and 20 RAND_20_-REG_10_. 20% of the sequences contained a gap, with equal probability spread across conditions. Gap lengths were as in Experiment 1B.

#### Experiment 3 Stimuli and Procedure

The stimulus set was identical to that in Experiment 1 but that stimuli contained no gaps. Participants were instructed to press a button as quickly as possible after detecting a pattern change in the sound sequence. Key presses were checked after each tone, so the resolution of the reaction time measurement was 50ms. In total 150 stimuli were presented in random order over 5 consecutive blocks – 25 of each condition –with an inter-trial interval of five to seven seconds.

#### Experiment 4A Stimuli and Procedure

The stimulus set consisted of the conditions used in Experiment 1 (Figure 1) and additionally included a new condition: **RAND_20_-REG_1_** – which consisted of a transition from a random sequence to a sequence of fixed frequency tones (REG_1_) (see Figure 7). In total, 288 trials were presented in random order over 8 consecutive blocks: 24 RAND_20_-REG_1_, 24 RAND_20_-REG_10_, 48 RAND_20_, 48 REG_10_, 48 REG_10_-RAND_20_, 48 CONT and 48 STEP, with one-third of the sequences containing a gap (lengths as in Experiment 1B).

#### Experiment 4B Stimuli and Procedure

The stimulus set was expanded to include two additional conditions: **RAND_20_-REG_2_** consisted of a transition from a random sequence to a sequence to a regular pattern of two alternating tones; **RAND_20_-REG_5_** consisted of a transition from a random sequence to a sequence to a regular pattern of five alternating tones (Figure 7). Overall, 168 stimuli were presented in random order over 7 consecutive blocks including 21 RAND_20_-REG_1_, 21 RAND_20_-REG_2_, 21 RAND_20_-REG_5_, 21 RAND_20_-REG_10_ and 84 RAND_20_, with one-third of the sequences containing a gap.

### QUANTIFICATION AND STATISTICAL ANALYSIS

#### Comparison of PDRs across conditions

A series of paired t-tests were conducted on each pair of conditions (two-tailed; over the entire epoch length; downsampled to 20 Hz), with family-wise error (FWE) control using a non-parametric permutation procedure with 5,000 iterations (cluster-defining height threshold of p <0.05 with an FWE-corrected cluster size threshold of p <0.05; Maris and Oostenveld, 2007), as implemented in the Fieldtrip toolbox (http://www.fieldtriptoolbox.org; Oostenveld et al., 2010). Significant time intervals are presented as colored horizontal bars below the PDR plots.

#### Event rate analysis

Because PD/PC events are rare (normal pre-transition rates are 1∼2 per second) the statistical analysis was conducted by pooling over Experiment 1A and B (32 subjects overall). The cluster analysis used to compare PDR was conducted between each transition condition and its control with other details as described above.

#### Experiment 3

To quantify the change in PDR peak latency for STEP and REG_10_-RAND_20_ in Experiment 3 (active transition detection) relative to Experiments 1,2,4,5 (gap detection), bootstrap analysis (1000 iterations; Efron and Tibshirani, 1994) was performed on two sets of participant data, one constructed from the 14 active participants in Experiment 3 (‘active’) and another from the 57 participants pooled from Experiments 1,2,4,5 (‘non-active’). On each iteration, a ‘simulated’ PDR latency was computed over 14 participants randomly drawn from the ‘non-active’ pool. The scatterplots in Figure 5C show the distribution of the simulated peak latency of STEP (left) and REG_10_-RAND_20_ (right) in the ‘non-active’ pool. The red crosses indicate the mean peak latency in Experiment 3.

We analyzed potential single-trial level associations between reaction time and pupil responses using REML in JMP 13.2 (SAS Institute, SAS Institute Inc., Cary, NC). Because reaction times (RTs) and pupillometry measures were non-normally distributed, rank scores were used for all analyses (ties assigned the bottom rank from the set of same values). Three measures of PDR dynamics were investigated: ‘**Max PDR amplitude’**, reflected the peak pupil diameter; **‘Max PDR latency’,** reflected the latency of the PDR peak**, ‘max derivative PDR latency’** measured the time of maximum pupil rate of change. Participant was entered as the random factor, and rank RT as fixed effect; for reported analyses, slope was fixed over subjects to avoid potential overfitting, but all effects at p < 0.05 also hold when separate slopes are fit for each participant (except when noted). We report t- and p-values associated with the RT fixed effect; we also provide an estimate of relative contribution of the fixed effect to overall model fit by computing partial R^2^ estimates using the lme4 and r2glmm (Jaeger et al., 2017) packages in R; these provide an implementation of the Nakagawa and Schielzeth (2013) algorithm.

### Data Availability

Data will be made freely available to readers from the date of publication. Specific details of the relevant repository will be provided upon manuscript acceptance.

## Supplementary Materials

**Figure S1:**
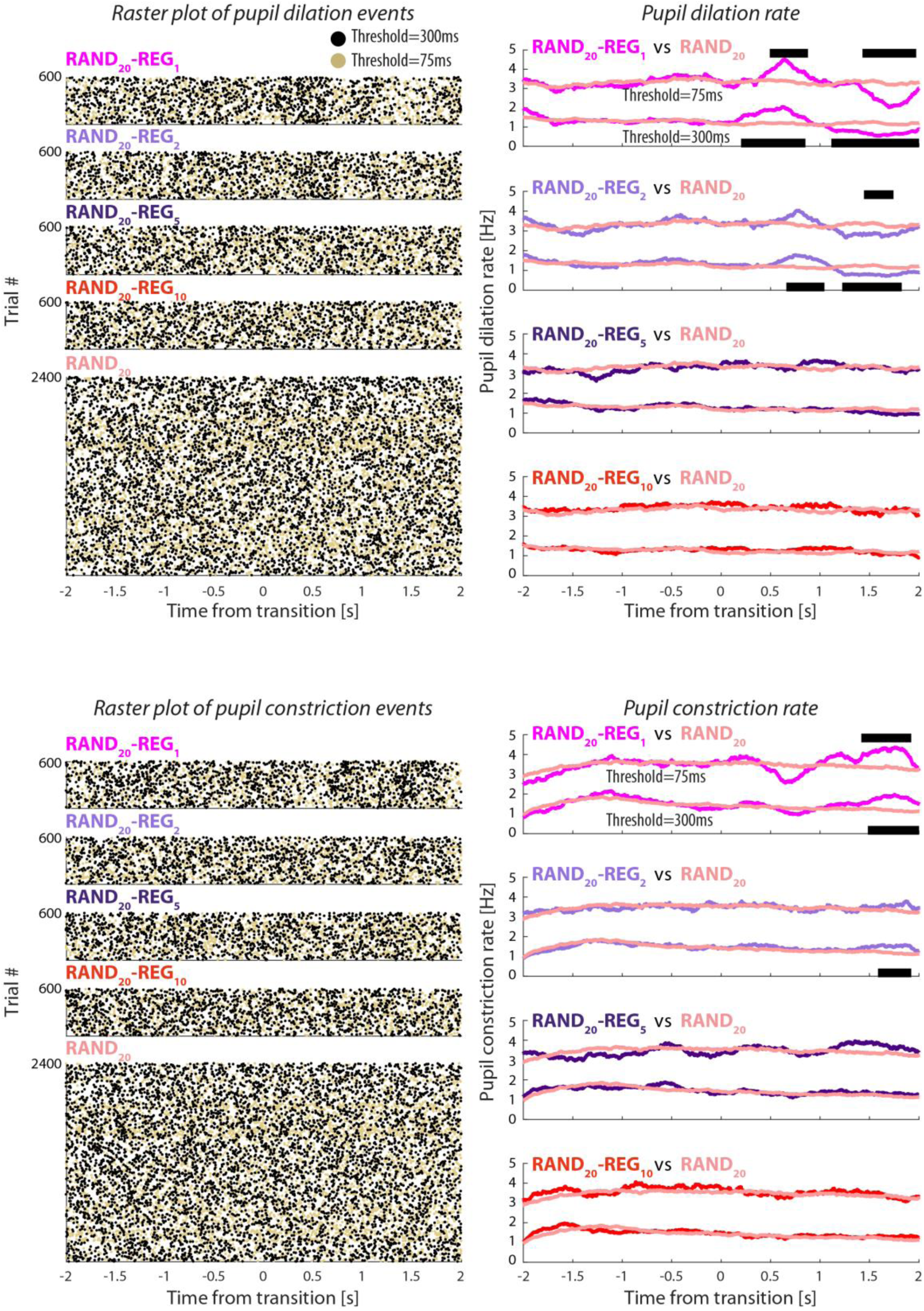
Experiment 4B: Pupil dilation and constriction rates. [Top left] Raster plots of pupil dilation (PD) events extracted from all trials and all participants (collapsed over the two groups, N = 30). Each line represents a single trial. Black dots represent the onset of a pupil dilation with a duration of at least 300ms, yellow dots represent pupil dilation onsets with a threshold duration of 75ms. Transition time is indicated by a black vertical line. [Top right] Pupil dilation rate (running average with a 500ms window) as a function of time relative to the transition. From top to bottom, one of four transition conditions are plotted against the no-change control RAND_20_: RAND_20_-REG_1_, RAND_20_-REG_2_, RAND_20_-REG_5_ and RAND_20_-REG_10_. The black horizontal lines indicate time intervals where cluster-level statistics showed significant differences between each change condition and the no-change control RAND_20_. The statistics for the PD rates with a threshold duration of 75ms and 300ms are placed above and below the graph, respectively. The lower panels present the pupil constriction (PC) rate results, arranged in the same format.

